# The rural urban divide, insights from immuno-genetic profiles and implications for health

**DOI:** 10.1101/2023.09.08.556879

**Authors:** Reem Hamad, Nahid A Eid, Safa A.E Aboswar, Azza Saeed, Ayman A Hussein, Ibrahim M Elhassan, Kirk A Rockett, Dominic P Kwiatkowski, Muntaser E Ibrahim, Ahmed M Elhassan

## Abstract

Population disparities in health and disease, has been observed, and amply documented. While often attributable to genetic underpinnings, such disparities, extends beyond population genetic predisposition to include environmental and geographic determinants most pronouncedly the division between rural and urban lifestyles. Under such influences, genes and gene products may become affected by epigenetic factors, microbial modifiers including infections and the body microbiome that ultimately shapes the outcome of complex milieu of protein networks. Retrospective, demographic, genotype, and expression data of two rural populations from eastern Sudan were analysed for genotype, allele frequency distribution, Hardy Weinberg Equilibrium and expression profiles using an array panel of Th1, Th2 and Th3 genes in a subset of rural population sample against matched urban control.

Differences between urban and rural samples were observed in the departure from HWE with excess of heterozygosity in the rural sample. In the Th1, Th2 and Th3 array, cytokines were consistently overexpressed in the rural as compared to urban cohort and was replicated in 7 selected genes that are associated with chronic diseases amongst urban dwellers in contrast to rural village inhabitants. IgE levels as feature of parasitic infections is one other difference to include in that dichotomy.

Gene expression appears to be more exposing to an overall outcome of genetic variations, including the interaction with environmental influences within and outside the body. Here, it may be gathered from the contrast in the expression patterns between the rural and urban samples. The presence of signals of natural selection in genes that are key to certain biological functions as CD40L and FasL, and the sharp contrast between urban and rural populations in gene variants distribution and expression patterns may provide important clues towards understanding the disparity between human communities in non-communicable diseases of lifestyle as well as some of the emerging infectious diseases.

## Introduction

Rural urban divides are among the earliest social structures in human history, with the term “civilization” often denoting that shift from rural based economies and habits into cities and urbanized lifestyles. Since the last glacial maxima, the process has been ensuing progressively reaching its zenith in the past few centuries with the industrial revolution. Such fundamental shift has its impact on the human biological and psychological well-being and disease transmission patterns. It has been speculated that a major trade-off took place whereby infectious diseases like Malaria and Helminthes i.e., communicable diseases prevalent in rural communities has been replaced by chronic and non-communicable diseases in urban environments with some far-reaching implications.

During its biological history, the human species were under continuous challenge by infectious agents.

It is the host-parasite interactions that shaped our evolutionary history particularly diseases like malaria which remains one of the major causes of mortality among children worldwide and thus among the strongest known forces for evolutionary selection in recent history (1). Analysis of malaria candidate genes from malaria endemic areas have indicated the potential impact of the population structure on the outcome of association in susceptibility to malaria and that the structure is apparently a function of both the unified ethnicity and relatedness within the population (2). The expression of these genes however is not necessarily a direct reflection of such variation and could be subject to the complex interaction of the protein networks of the body and that with the environment.

In the rapidly escalating urban centres, socioeconomic differences result in diverse environmental acquaintances which largely determine disease patterns. Understanding the factors associated with altering environments and the link to the change of the immune system would be important for both communicable and non-communicable diseases prevention strategies. Although risk factors associated with the rural to urban transition, particularly relative to inflammatory diseases, have been studied comprehensively (3,4) little is known though about changes that takes place in the immune system as function of the rural–urban gradient.

During routine longitudinal health surveillances of rural population in eastern Sudan (2), we have noticed a striking dearth of chronic illnesses like diabetes, hypertension, cancer and Asthma, while these illnesses are registering steady increase in urban populations nationwide. Taken together this shift forms what is known as the health features of the transition, the term coined in reference to the changes from rural mostly subsistent economy into an urbanized based lifestyle (reference, article on the socio biological). The association of the transition with diseases like Asthma has been amply described, however no proper studies on the underlying genomic and molecular basis of chronic illnesses in relation to transition has been carried out.

Despite the bountiful of studies that tackled the genetic elements of gene expression, few have addressed the geographical location impact at transcriptional level. Idaghdour and his colleagues studied the Amazighs of Morocco -a relatively genetically homogenous population-that lives in three distinct geographical localities, they concluded that geographical area differences might have a dominant effect on gene expression profiles up to one third of the leukocyte transcriptome (5,6). Investigating gene expression patterns is one approach toward scrutinising differences in immune responses between urban and rural populations, as variability in gene expression is a result of environmental factors in addition to genetic ones (7). Moreover, differential gene expression can be a key mechanism in disease manifestation. (8).

To date, a lack of data has hampered the identification of functional genomic features that fosters these differences, that might have led to better understanding as well as discovery of specific roles of genes in immunity. Here we investigate whether contrasting cultural and geographical locations in urban and rural areas in Sudan that are endemic to malaria and leishmaniasis may have had an impact on mRNA expression of selected immune genes among population and consequently their health and disease profiles.

## Materials and Methods

### Ethics Statement

Analysis based on archived data and research projects approved by the Ethical Committee of the Institute of Endemic Diseases, University of Khartoum, was carried out. Samples were originally taken with written informed consent from all individuals.

### Study population and sample size

Genotype and phenotype data were analysed retrospectively from two Rural village populations of the Hausa and Massalit ethnicities. These populations reside in the malaria-endemic area of the Rahad River in Eastern Sudan and have been under routine surveillance longitudinally over the past decades for different infectious diseases including malaria and leishmaniasis (2).

A total of 168 SNPs were genotyped in 414 individuals from Hausa and 510 from Masalit using the Sequenom® iPLEX gold assay. The SNPs were originally chosen based on previously published reports of malaria candidate-gene associations in addition to SNPs that has shown early promise for associations in a Genome Wide Study undertaken in the Gambia by the MalariaGEN consortium. No urban controls were genotyped in the MalariaGEN project but controls consisting of 60 supposedly immunologically naïve samples for infectious diseases, were selected based on an extended urban lifestyle and habitation were included, of those 20 samples had their RNA extracted and subsequently analyzed for gene expression as below.

### Expression analysis

To estimate the gene expression profile, we used the RT2 Profiler PCR Array for Th1/Th2/Th3 panel 96 set of primers that include 84 cytokine genes involved in immune response, to analyse the expression profile of the samples from Rural and Urban areas we used an online program in the web site http://www.sabiosciences.com/rt_pcr_product/HTML/PAHS-011A.html of the company. Then, 7 genes were selected for further expression analysis in the rural and urban areas using Rt-PCR based on their putative association with chronic diseases including cancer and their signalling pathways. For the single gene expression, the following formula was used:

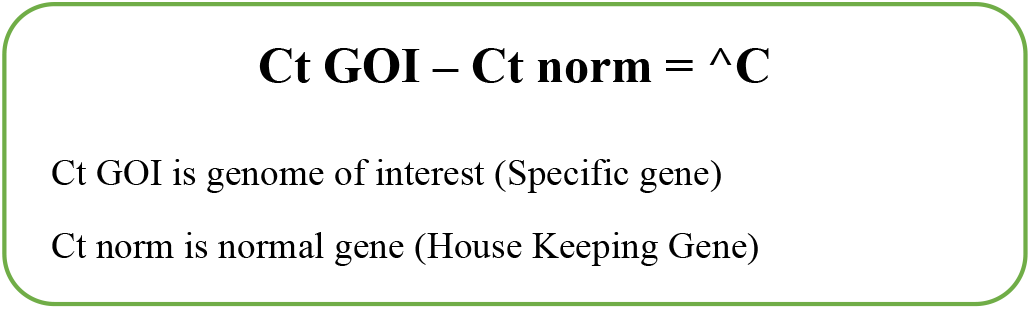

### Immunoglobulin E(IgE) level Estimation

Stored blood samples collected in previous cross-sectional surveys in the years 2000,2001, 2005, 2007 from two village populations were analysed, 50-60 subjects for IgE level using ELISA.

## Results

Helminthic infections are also prevalent but their incidence in these rural communities are not properly determined except for a limited survey in school children (157 in Um Salala and 44 in KoKa), where in Koka only 7 cases (5 cases of Taenia, 1 case of H-nana and one of Enterobiasis), while in Salala there were 39 cases (1 case of Ascaris, 8 cases of H-nana, 24 cases of Taenia and 6 cases of Schistosoma mansoni).

### Immunoglobulin E (IgE) level

The normal value range of IgE is 0-380 IU/ml, both villages show yearly fluctuation in the average IgE level which varied between the two villages with Koka village consistently having higher readings than Salala. This variability was not significant in the cross-sectional survey of years 2000 and 2001. However, in the years, 2005, 2007 the readings in Koka were much higher than the normal values being 710, 723 IU/ml in the two years respectively while in Salala the level was 387,374 IU/ml, a significant difference with *P* ≤ 0.00081. (Fig 1).

**Fig 1:**
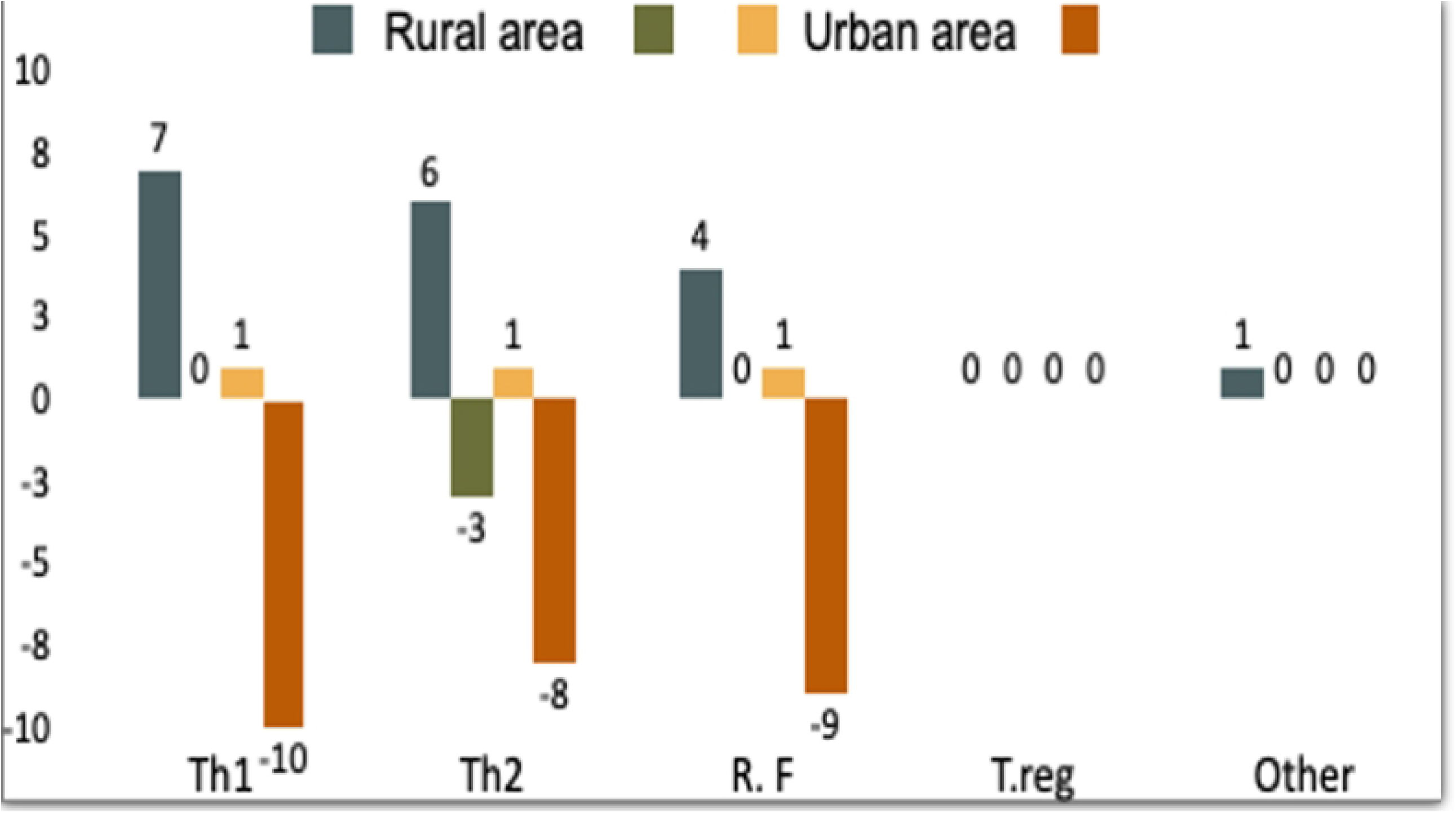
Average IgE level in Koka and Salala in year 2000,2001,2005and 2007

### Genotype analysis

Malaria-Gen genotype data originally generated for association studies on genetic susceptibility to malaria were used here to capture signals of natural selection by infectious diseases in rural population, and as a likely explanation for the differences in the expression patterns.

Departure from HWE was observed in 14 SNPs in both cases and controls (Table1&Ttable 2) indicating either impact of natural selection or effects of population sub structuring. A control sample from the genome database which is predominantly urban was analysed and the results shows only 4 of those SNPs are in departure from HWE.

**Table 1:** Hardy-Weinberg Equilibrium and significant association P value between urban and rural (Massalit)

|  | HWE_P_Case | HWE_P_Control | Urban sample |
| --- | --- | --- | --- |
| **LOC44rs2706381** | NS | NS | NS |
| **LOC441108.rs7704457** | **0.00284** | **0.6426** | NS |
| **IL3.rs35415145** | NS | NS | 0.014851485 |
| **IL5.rs2069818** | NS | NS | NS |
| **IL22 rs2227485** | **0.83832** | **0.020623** | NS |
| **IL22 rs2227478** | **0.82505** | **0.024136** | NS |
| **IL1A rs17411697** | **0.56827** | **0.01637** | NS |
| **CTL4 rs2242665** | NS | NS | NS |
| **IL22 rs2227491** | **0.72206** | **0.005468** | NS |
| **IL22 rs1012356** | **0.13729** | **0.039258** | **NS** |
| **RAD rs4621555** | **0.00347** | **0.66165** | **NS** |
| **IRF1.rs2070729** | **NS** | **NS** | **NS** |
| **IL13.rs848** | **NS** | **NS** | **NS** |
| **CD40LG rs3092945** | **0.03279** | **7.15E-11** | **NS** |
| **G6PD-376.rs1050829** | **NS** | **NS** | **NS** |
| **IL17RE.rs708567** | **NS** | **NS** | **NS** |
| **IL4.rs2243283** | **NS** | **NS** | **NS** |
| **IL4 rs2243270** | **0.28887** | **0.02693** | **NS** |
| **IL20RA rs1555498** | **0.92658** | **0.04570** | **NS** |
| **ABO rs8176719** | **0.52372** | **0.00126** | **NS** |
| **P4HA2 rs3805685** | **0.14016** | **0.00159** | **0.044554455** |

NS: not significant

**Table 2:** Hardy-Weinberg Equilibrium and significant association P value between Urban and Rural(Hausa).

|  | **HWE_P_Case** | **HWE_P_Control** | **Urban sample** |
| --- | --- | --- | --- |
| **LOC44rs2706381** | **0.02613** | **0.018582** | **NS** |
| **LTA rs909253** | **0.88956** | **0.0068871** | **0.04950495** |
| **LTA rs2239704** | **NS** | **NS** | **NS** |
| **IL5.rs2069818** | **NS** | **NS** | **NS** |
| **IL22 rs2227485** | **0.048194** | **0.45624** | **NS** |
| **IL22 rs2227478** | **0.028801** | **0.88961** | **NS** |
| **CD36 rs3211938** | **0.3396** | **0.04037** | **0.014851485** |
| **CTL4 rs2242665** | **NS** | **NS** | **NS** |
| **RTN3 rs542998** | **0.77931** | **0.048729** | **NS** |
| **GNAS rs8386** | **NS** | **NS** | **NS** |
| **DERL3.rs3177244** | **NS** | **NS** | **NS** |
| **CFTR rs17140229** | **NS** | **NS** | **NS** |
| **ADORA2B rs2535611** | **0.94521** | **0.041863** | **NS** |
| **CD40LG rs3092945** | **6.70E-06** | **5.31E-09** | **NS** |
| **CD40LG.rs17424229** | **0.097609** | **1.93E-08** | **NS** |

### Th1/Th2/Th3 Array expression profile

There were clear differences in the pattern of expression between rural and urban areas. Overall, up regulated genes were dominant in the rural), while down regulation was the predominant mode in the urban samples. Th2 samples were down regulated in several rural samples (Fig 2). Number of genes down or upregulated were indicated in each column which represents a single sample.

**Fig 2:**
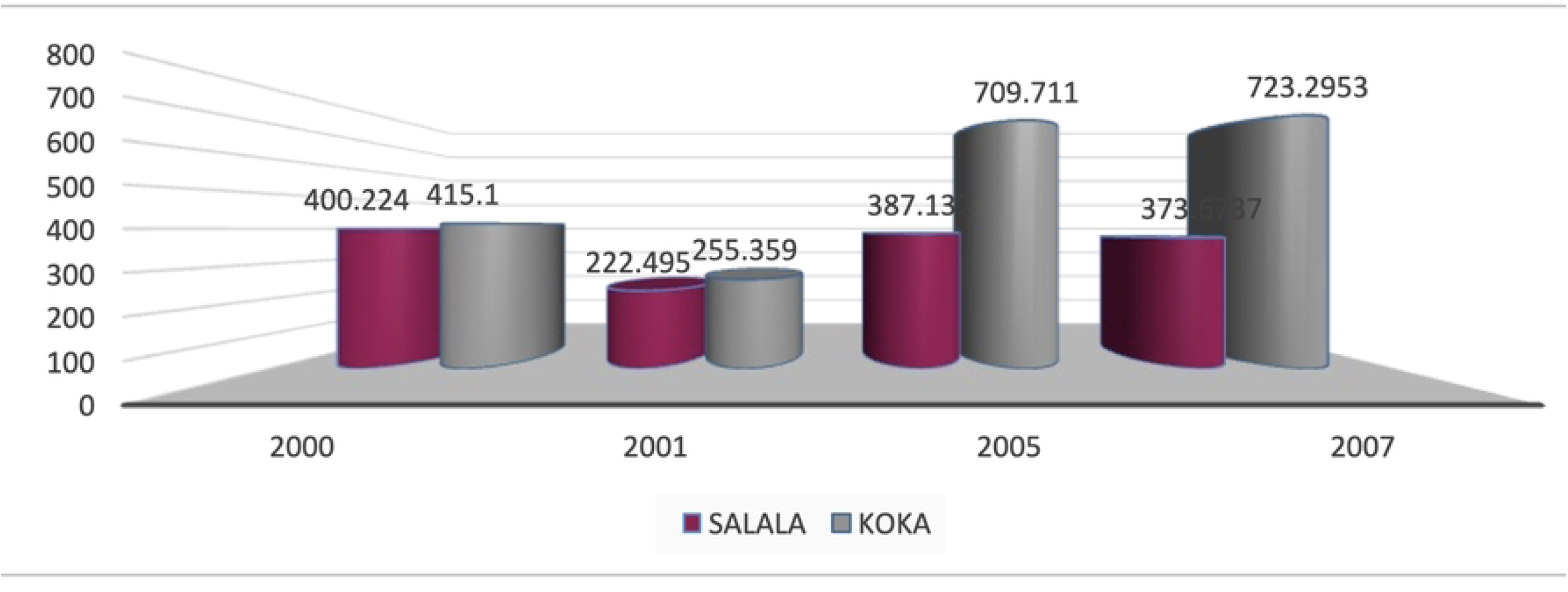
Expression profile for cytokine genes (Th1, Th2, Th3 array), differences as measured against a control point from each rural and Urban areas. There are clear differences in the pattern of expression between rural and urban areas. Overall, up regulated genes were dominant in the rural), while down regulation was the predominant mode in the urban samples. Th2 samples were down regulated in several rural samples.

### Gene expression profile

The analysis of the 10 genes were selected from those expressed through all samples. Seven genes were analysed for both areas. Again, down regulation of expression of the selected genes was the main feature among the urban (n= 23) in comparison to rural area (Salala n= 21, Koka, n=29).

Unlike the array expression few rural samples showed down expression tendency and three urban sample showed slight over expression in one gene (CTLA4). (Fig 3)

**Fig 3:**
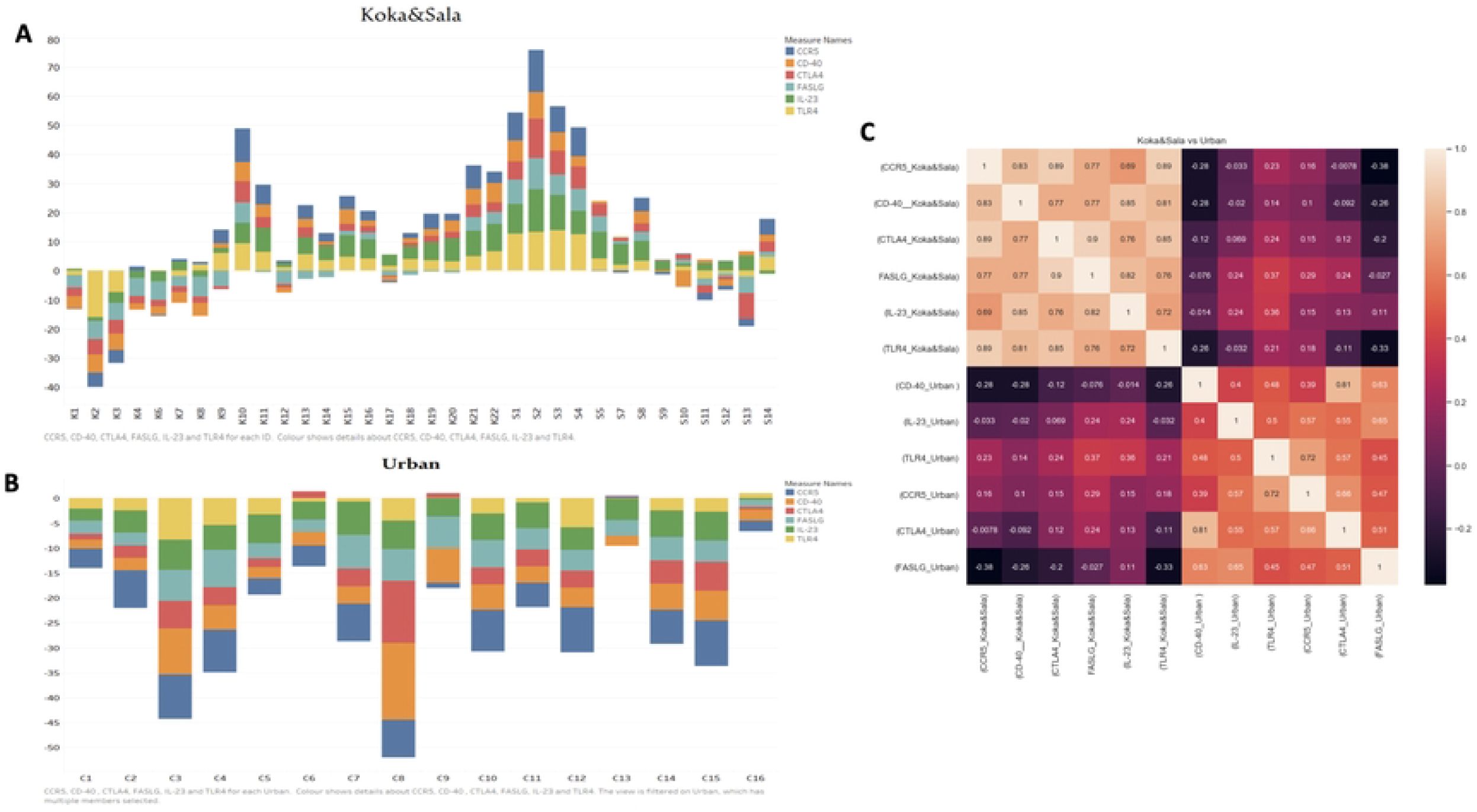
Shows Selected gene expression difference between Urban and rural (Koka and Salala). A. Rural expression bar chart. B. Urban expression Bar chart. C. Heat map of combined urban and rural expression.

## Discussion

In the human host, the outcome of infectious disease is determined by a complex interactive relationship between the host, the parasite, and the environment. Against such complex multifactorial milieu, the dichotomous health disparity between rural and urban lives are considered. This obvious, dichotomy has been validated by various reports and anecdotes across the globe, particularly in developing countries (9-13). Underlying these disparities are multiple elements involving social and environmental factors that characterize and distinguish urban and rural spaces, and thereby affect health profiles and risk of contracting diseases. Despite observations and ample documentation, few studies attempted to venture understanding the genotypic and immunological differences in expression between the two spaces. In one setting humans are in tandem with their evolutionary history, in terms of nutrition, exposure to pathogens and psychosocial elements like crowding and nature of household etc. On the other hand, people are subject to new nutritional cultures, pathogens free environment and stress where the repercussions on human health are enormous. Among the interesting observations that may provide some clues onto this dichotomous health status, are the expression profiles of candidate genes originally tested within the Malaria-Gen consortium of which some were included in the Th1/Th2/Th3 arrays. Several of these genes are candidates for disparate burden of chronic diseases like Asthma (IL-.4, IL-5 and TLR4) and cancer (CD-40, FASL, CTLA4 and CCR5) and are being investigated against the reported extremely low incidence of Asthma and cancer in the area.

Rural-urban differences seen in the current data sets, were of great interest. Despite the small urban sample size, in the overall outcome, both communities corresponded to the dichotomous urban-rural divide. How much natural selection is responsible for such differences in expression remains to be answered. Populations of the rural area sampled in this study were a subject of extended investigation over three decades for transmission of infectious diseases where the area is endemic for both malaria and visceral leishmaniasis (3,14,15). Conversely prolonged exposure overtime in some cases may result in genetic and epigenetic differences. In these villages the impact of natural selection in some candidate genes is possibly due to pressure by infectious diseases like VL rather than malaria (16). Natural selection is best seen in the departure from Hardy Weinberg expectations which is shown in a group of loci including CTLA4 and CFTR. Interestingly, it is recorded more frequently in controls than cases suggesting that these signals might emanate from reasons other, in addition to natural selection, like sampling effects and population structure both known causes of DHWE to be considered. Whether helminthic infections could explain some of the health features in these rural communities including elevated IgE a Bonafede marker of Asthma remain to be established. Bearing in mind that IgE has been seen to be elevated during malaria infection (17), differences in IgE levels between the two villages, however, could not be ascribed solely to malaria as the prevalence and incidence rates of the disease are quite comparable between the villages (2). Such an observation might insinuate an innate/genetic component in the Hausa community that favours the production of high IgE levels. In Nigeria, a study was done among children where the mean serum level of IgE was significantly raised in children with helminthic or protozoan infection. In contrast, the level of IgE was reduced in children with *Ascaris lumbricoides* compared to Plasmodium species (18). While another study in Ghana found elevated level of IgE among rural inhabitants explained by high incidence of helminthic infections (19).

Differences in malaria transmission between rural and urban centres (20), could be one of the main drivers of these patterns, as several of these markers like IL-4, IL-5, IL-10, and IL-13 are markers of CD4 T-Helper 2 activity (21). Lower CD40 activity in urban compared to rural populations, conforms to the literature of higher susceptibility to chronic diseases among urban populations (22). On the other hand, that low burden of chronic diseases like Asthma, diabetes, and cancer in rural communities, could have complex underlying molecular aetiologies beside lifestyle.

However, it might prove challenging to pinpoint or specify actual effector molecule(s) in that trade-off between chronic and infectious diseases, given the intricacies of signalling pathways and epigenetics determinants. One might be tempted, though, to hypothesize that the observed low cancer incidence in these villages might have something to do with the endemicity of the malaria parasite. Salanti et al., (2015), have shown homology between cancer cells and malaria parasite and hence antibodies against malaria protein may be potentially protective against cancer (23). TNF is obviously another potential candidate for these trade-offs. In Ghana, it was found that TNF-a, IL-6, and IL-12 levels in urban dwellers with malaria were significantly higher, while IL-10, CD4, CD3, CD8T-cells levels and CD4/ CD3 ratio were significantly lower (24). However, in the current dataset, the urban population displayed an overall low expression of key cytokine genes in both Th1/Th2 classes which is consistent with lower exposure to microbial environment, something observed in some other studies (25). In comparing the expression between three geographical distinct groups in Indonesia a significant association was found especially among genes involved in immune function suggesting a potentially adaptive response to local environmental pressures (26).

One limitation of the current study is being conducted with relatively small numbers of subjects in each study area. A larger sample size with several points of collection over time, may have meant greater statistical power in detecting area differences for some of the genes. Common to all mRNA studies of whole blood, our study also suffers from the fact that mRNA expression might not be directly related to protein expression levels. In addition, the expression of the mRNA is in whole blood and does not reveal any cell-specific profiles which might be important when considering their function in determining disease profiles.

Nevertheless, results and observation reported here addresses some of the crucial matters pertaining to public health specially in developing countries under changing disease patterns accompanying change of lifestyle. Given that subsistence economies in rural Africa like in the Gedaref state are vanishing modes of life, leaves us with a narrow window of opportunity to foster an in-depth knowledge onto the biology of health transition that might inform future health policies and research directions.

## Acknowledgments

Authors are indebted to the inhabitants of the villages of Koka and Salala who diligently cooperated in providing biological samples to continued health surveillance in the Rahad River locality. Thus, making it possible to unlock much of the peculiarities of the disease’s endemic to their area. In turn the manuscript is dedicated to the memories of two giant figures in science, who co-authored this manuscript before passing away: A M Elhassan and D P Kwiatkowski whose tribute to the well-being of populations of these villages remains some of their everlasting legacy.

## References

1 Kwiatkowski DP. How malaria has affected the human genome and what human genetics can teach us about malaria. Am J Hum Genet. 2005 Aug;77(2):171–92. doi: 10.1086/432519. Epub 2005 Jul 6. PMID: 16001361; PMCID: PMC1224522.

2 Eid, N. a, Hussein, A. a, Elzein, A. M., Mohamed, H. S., Rockett, K. a, Kwiatkowski, D. P., & Ibrahim, M. E. (2010). Candidate malaria susceptibility/protective SNPs in hospital and population-based studies: the effect of sub-structuring. Malaria journal, 9, 119. doi:10.1186/1475-2875-9-119.

3 Fezeu L, Balkau B, Kengne AP, Sobngwi E, Mbanya JC. Metabolic syndrome in a sub-Saharan African setting: central obesity may be the key determinant. Atherosclerosis. 2007;193(1):70–76. doi:10.1016/j.atherosclerosis.2006.08.037

4 Delisle H, Ntandou-Bouzitou G, Agueh V, Sodjinou R, Fayomi B. Urbanisation, nutrition transition and cardiometabolic risk: the Benin study. Br J Nutr. 2012;107(10):1534–1544. doi:10.1017/S0007114511004661

5 Idaghdour Y, Storey JD, Jadallah SJ, Gibson G. A genome-wide gene expression signature of environmental geography in leukocytes of Moroccan Amazighs. PLoS Genet. 2008;4(4):e1000052. Published 2008 Apr 11. doi:10.1371/journal.pgen.1000052

6 Idaghdour Y, Czika W, Shianna KV, et al. Geographical genomics of human leukocyte gene expression variation in southern Morocco. Nat Genet. 2010;42(1):62–67. doi:10.1038/ng.495

7 Zeller T, Wild P, Szymczak S, et al. Genetics and beyond--the transcriptome of human monocytes and disease susceptibility. PLoS One. 2010;5(5):e10693. Published 2010 May 18. doi:10.1371/journal.pone.0010693

8 Cookson W, Liang L, Abecasis G, Moffatt M, Lathrop M. Mapping complex disease traits with global gene expression. Nat Rev Genet. 2009;10(3):184–194. doi:10.1038/nrg2537

9 Abdesslam B. Evolution of rural-urban health gaps in Morocco: 1992-2011. BMC Res Notes. 2012;5:381. Published 2012 Jul 27. doi:10.1186/1756-0500-5-381

10 Fotso JC. Urban-rural differentials in child malnutrition: trends and socioeconomic correlates in sub-Saharan Africa. Health Place. 2007;13(1):205–223. doi:10.1016/j.healthplace.2006.01.004

11 Fotso JC. Child health inequities in developing countries: differences across urban and rural areas. Int J Equity Health. 2006;5:9. Published 2006 Jul 11. doi:10.1186/1475-9276-5-9

12 Fox K, Heaton TB. Child nutritional status by rural/urban residence: a cross-national analysis. J Rural Health. 2012;28(4):380–391. doi:10.1111/j.1748-0361.2012.00408.x

13 Liu H, Fang H, Zhao Z. Urban-rural disparities of child health and nutritional status in China from 1989 to 2006. Econ Hum Biol. 2013;11(3):294–309. doi:10.1016/j.ehb.2012.04.010

14 Salih NA, Hussain AA, Almugtaba IA, et al. Loss of balancing selection in the betaS globin locus. BMC Med Genet. 2010;11:21. Published 2010 Feb 3. doi:10.1186/1471-2350-11-21

15 Himeidan, Y. E., Elzaki, M. M., Kweka, E. J., Ibrahim, M., & Elhassan, I. M. Pattern of malaria transmission along the Rahad River basin, Eastern Sudan. Parasites & vectors. 2011; 4: 109. doi:10.1186/1756-3305-4-109.

16 Elhassan AA, Hussein AA, Mohamed HS, et al. The 5q31 region in two African populations as a facet of natural selection by infectious diseases. Genetika. 2013;49(2):279–288. doi:10.7868/s0016675813020057.

17 Elghazali G, Perlmann H, Rutta AS, Perlmann P, Troye-Blomberg M. Elevated plasma levels of IgE in Plasmodium falciparum-primed individuals reflect an increased ratio of IL-4 to interferon-gamma (IFN-gamma)-producing cells. Clin Exp Immunol. 1997;109(1):84–89. doi:10.1046/j.1365-2249.1997.4401337.x

18 Winter WE, Hardt NS, Fuhrman S. Immunoglobulin E: importance in parasitic infections and hypersensitivity responses. Arch Pathol Lab Med. 2000;124(9):1382–1385. doi:10.5858/2000-124-1382-IE

19 Amoah AS, Obeng BB, May L, et al. Urban-rural differences in the gene expression profiles of Ghanaian children. Genes Immun. 2014;15(5):313–319. doi:10.1038/gene.2014.21

20 Robert V, Macintyre K, Keating J, et al. Malaria transmission in urban sub-Saharan Africa. Am J Trop Med Hyg. 2003;68(2):169–176.

21 Zhu J, Paul WE. CD4 T cells: fates, functions, and faults. Blood. 2008;112(5):1557–1569. doi:10.1182/blood-2008-05-078154

22 Acheampong DO, Adu P, Ampomah P, Duedu KO, Aninagyei E. Immunological, haematological, and clinical attributes of rural and urban malaria: a case-control study in Ghana. J Parasit Dis. 2021;45(3):806–816. doi:10.1007/s12639-021-01363-4

23 Salanti A, Clausen TM, Agerbæk MØ, et al. Targeting Human Cancer by a Glycosaminoglycan Binding Malaria Protein. Cancer Cell. 2015;28(4):500–514. doi:10.1016/j.ccell.2015.09.003

24 Acheampong DO, Adu P, Ampomah P, Duedu KO, Aninagyei E. Immunological, haematological, and clinical attributes of rural and urban malaria: a case-control study in Ghana. J Parasit Dis. 2021;45(3):806–816. doi:10.1007/s12639-021-01363-4

25 M’Bondoukwé NP, Moutongo R, Gbédandé K, et al. Circulating IL-6, IL-10, and TNF-alpha and IL-10/IL-6 and IL-10/TNF-alpha ratio profiles of polyparasitized individuals in rural and urban areas of gabon. PLoS Negl Trop Dis. 2022;16(4):e0010308. Published 2022 Apr 14. doi:10.1371/journal.pntd.0010308

26 Natri HM, Bobowik KS, Kusuma P, et al. Genome-wide DNA methylation and gene expression patterns reflect genetic ancestry and environmental differences across the Indonesian archipelago. PLoS Genet. 2020;16(5):e1008749. Published 2020 May 26. doi:10.1371/journal.pgen.1008749

